# Can we predict real-time fMRI neurofeedback learning success from pre-training brain activity?

**DOI:** 10.1101/2020.01.15.906388

**Authors:** Amelie Haugg, Ronald Sladky, Stavros Skouras, Amalia McDonald, Cameron Craddock, Matthias Kirschner, Marcus Herdener, Yury Koush, Marina Papoutsi, Jackob N. Keynan, Talma Hendler, Kathrin Cohen Kadosh, Catharina Zich, Jeff MacInnes, Alison Adcock, Kathryn Dickerson, Nan-Kuei Chen, Kymberly Young, Jerzy Bodurka, Shuxia Yao, Benjamin Becker, Tibor Auer, Renate Schweizer, Gustavo Pamplona, Kirsten Emmert, Sven Haller, Dimitri Van De Ville, Maria-Laura Blefari, Dong-Youl Kim, Jong-Hwan Lee, Theo Marins, Megumi Fukuda, Bettina Sorger, Tabea Kamp, Sook-Lei Liew, Ralf Veit, Maartje Spetter, Nikolaus Weiskopf, Frank Scharnowski

## Abstract

Neurofeedback training has been shown to influence behavior in healthy participants as well as to alleviate clinical symptoms in neurological, psychosomatic, and psychiatric patient populations. However, many real-time fMRI neurofeedback studies report large interindividual differences in learning success. The factors that cause this vast variability between participants remain unknown and their identification could enhance treatment success. Thus, here we employed a meta-analytic approach including data from 24 different neurofeedback studies with a total of 401 participants, including 140 patients, to determine whether levels of activity in target brain regions during pre-training functional localizer or no-feedback runs (i.e., self-regulation in the absence of neurofeedback) could predict neurofeedback learning success. We observed a slightly positive correlation between pre-training activity levels during a functional localizer run and neurofeedback learning success, but we were not able to identify common brain-based success predictors across our diverse cohort of studies. Therefore, advances need to be made in finding robust models and measures of general neurofeedback learning, and in increasing the current study database to allow for investigating further factors that might influence neurofeedback learning.

## Introduction

During the last years, neurofeedback using real-time functional magnetic resonance imaging (fMRI) has been gaining increasing attention in cognitive and clinical neuroscience. Real-time fMRI-based neurofeedback enables subjects to learn control over brain activity in localized regions of interest (ROIs). Brain areas that have been investigated in fMRI-based neurofeedback studies include the anterior cingulate cortex (deCharms et al., 2005; Emmert et al., 2014; Gröne et al., 2015; Guan et al., 2014; Li et al., 2013), anterior insula (Yao et al., 2016), amygdala (Brühl et al., 2014; Gerin et al., 2016; Keynan et al., 2016; Nicholson et al., 2017; Paret et al., 2014; Young et al., 2014), auditory cortex (Emmert, Kopel, Koush, Maire, Senn, Van De Ville, et al., 2017; Haller, Birbaumer, & Veit, 2010), default mode network (DMN; McDonald et al., 2017), dorsolateral prefrontal cortex (Sherwood, Kane, Weisend, & Parker, 2016), hippocampus (Skouras et al., 2019), insula (Korhan Buyukturkoglu et al., 2015; Caria et al., 2007; Emmert et al., 2014; Frank et al., 2012; Zilverstand, Sorger, Sarkheil, & Goebel, 2015), motor cortex (Auer, Schweizer, & Frahm, 2015; Blefari, Sulzer, Hepp-Reymond, Kollias, & Gassert, 2015; Buyukturkoglu et al., 2013; Marins et al., 2015; Scharnowski et al., 2015; Yoo, Lee, O’Leary, Panych, & Jolesz, 2008), nucleus accumbens (Greer, Trujillo, Glover, & Knutson, 2014), parahippocampal gyrus (Scharnowski et al., 2015), ventral tegmental area (MacInnes, Dickerson, Chen, & Adcock, 2016; Sulzer et al., 2013), and the visual cortex (Scharnowski, Hutton, Josephs, Weiskopf, & Rees, 2012; Shibata, Watanabe, Sasaki, & Kawato, 2011). More recently, functional brain networks have also been successfully trained employing connectivity-informed neurofeedback in networks sub-serving emotion regulation (Koush et al., 2015), attention (Koush et al., 2013), motor control (Liew et al., 2016; Megumi, Yamashita, Kawato, & Imamizu, 2015), craving (Kim, Yoo, Tegethoff, Meinlschmidt, & Lee, 2015), and executive control (Spetter et al., 2017).

Real-time fMRI neurofeedback has been shown to improve behavioral and cognitive functions in healthy participants (e.g. Rota et al., 2009; Scharnowski et al., 2015, 2012; Sherwood et al., 2016; Shibata et al., 2011), and to reduce clinical symptoms in neurological and psychiatric patient populations, such as patients suffering from adipositas (Frank et al., 2012), alcohol and nicotine addiction (Canterberry et al., 2013; Hanlon et al., 2013; Hartwell et al., 2016; Karch et al., 2015; Kim et al., 2015; Li et al., 2013), borderline personality disorder (Paret et al., 2016), chronic pain (deCharms et al., 2005; Guan et al., 2014), depression (Linden et al., 2012; Young et al., 2017, 2014), Huntington’s disease (Papoutsi et al., 2018), obsessive compulsory disorder (Buyukturkoglu et al., 2015), Parkinson’s disease (Buyukturkoglu et al., 2013; Subramanian et al., 2011), phobia (Zilverstand et al., 2015), post-traumatic stress disorder (Gerin et al., 2016; Nicholson et al., 2017), and tinnitus (Emmert, Kopel, Koush, Maire, Senn, Van De Ville, et al., 2017; Haller et al., 2010).

However, not every individual can benefit from neurofeedback training and neurofeedback learning success differs substantially between individuals. In fact, many studies report participants who were unable to gain control over their own brain activity, even after multiple training sessions. In these studies, an average of about 38% of all participants failed to modulate their own brain activity and were not able to reach predefined goals after neurofeedback training (Bray, Shimojo, & O’Doherty, 2007; Chiew, LaConte, & Graham, 2012; deCharms et al., 2005; Johnson et al., 2012; Ramot, Grossman, Friedman, & Malach, 2016; Robineau et al., 2014; Scharnowski et al., 2012; Yoo et al., 2008). This failure to modulate brain activity, also referred to as the “neurofeedback inefficacy problem” (Alkoby, Abu-Rmileh, Shriki, & Todder, 2017), leads to a reduction in overall efficiency of neurofeedback training and hampers translation to clinical interventions. To date, the factors that cause neurofeedback inefficacy as well as the large inter-individual variability in neurofeedback learning success in the field of real-time fMRI neurofeedback remain unknown.

Interestingly, neurofeedback studies based on electroencephalography (EEG) have reported very similar numbers of participants failing to gain control over their own brain activity (e.g. Enriquez-Geppert et al., 2014; Zoefel, Huster, & Herrmann, 2011). However, despite intrinsic similarities shared by neurofeedback tasks across imaging modalities, EEG-based and fMRI-based neurofeedback differ substantially with regards to the underlying technology, methods and mechanisms. In this meta-analysis, we focus selectively on fMRI-based neurofeedback; for an overview of successful predictors in EEG-based neurofeedback we refer interested readers to Alkoby et al. (Alkoby et al., 2017).

Here, we investigate the influence of neural activity before neurofeedback training on neurofeedback learning success. In particular, we ask whether activity levels in the neurofeedback target region(s) during pre-training no-feedback runs – runs where participants modulate their brain activity in the targeted ROI without neurofeedback – or functional localizer runs can predict neurofeedback learning success in subsequent neurofeedback training runs. As pre-training brain activity already contains information on factors such as participant compliance and responsiveness to specific stimuli, we hypothesized that specific signal features (e.g., brain activity levels) extracted from the trained ROI during no-feedback or localizer runs before neurofeedback training are correlated with the respective participant’s success in modulating their own brain activity. To test this hypothesis, we performed a meta-analysis on data from 24 real-time fMRI neurofeedback studies (see Table 1), including a range of different target brain areas (> 20 ROIs), participants (261 healthy participants and 140 patients), and neurofeedback training methods (activity-based feedback as well as connectivity-based feedback).

## Material and methods

### Included studies

This meta-analysis required data that cannot be extracted from publications alone. Therefore, we identified suitable real-time fMRI neurofeedback studies via the mailing list of the real-time functional neuroimaging community, and by directly contacting authors of real-time fMRI neurofeedback studies. Inclusion criteria were at least one task-based functional run engaging the trained ROI/ROIs prior to neurofeedback training (e.g. a functional localizer run, a no-feedback run, or a task engaging the target ROI that was not used for localization). For increased generalizability, we did not limit this study to a specific participant cohort, target ROI, or neurofeedback training method. All identified contributions were screened, and if data met the inclusion criteria, they were included in this meta-analysis (Moher et al., 2009).

In total, we included 24 independent neurofeedback studies with data from 261 healthy participants [studies 1,2, 4, 5, 7, 9 – 15, 18 – 21,24] and 140 patients, including patients with alcohol abuse or dependence [14], anxiety disorder [14], cannabis abuse [14], cocaine use disorder [7], depression [14, 23], Huntington’s disease [16, 17], obesity [22], obsessivecompulsive disorder [14], opioid abuse [14], schizophrenia [8], specific phobia [14], tinnitus [3], and tobacco use disorder [6]. 18 studies conducted neurofeedback training on brain activity, while another eight studies provided connectivity-based feedback (two studies investigated both activity- and connectivity-based neurofeedback). We did not receive data from studies that performed neurofeedback based on other measures, such as multivariate pattern analysis. Brain areas that were targeted in these studies include the amygdala, anterior cingulate cortex (ACC), anterior insula, auditory cortex, dorsolateral prefrontal cortex (dlPFC), dorsomedial prefrontal cortex (dmPFC), medial prefrontal cortex (mPFC), orbitofrontal cortex (OFC), parahippocampal gyrus (PHG), posterior cingulate cortex (PCC), precuneus, premotor cortex (PMC), primary motor cortex (M1), somatomotor cortex (SMC), superior parietal lobule (SPL), supplementary motor area (SMA), ventral tegmental area (VTA), ventromedial prefrontal cortex (vmPFC), and the visual cortex (Figure 1). Table 1 provides an overview over all studies (Auer et al., 2015; Blefari, Sulzer, Hepp-Reymond, Kollias, & Gassert, 2015b; Emmert, Kopel, Koush, Maire, Senn, De Ville, et al., 2017; Kim et al., 2015; Kirschner et al., 2018; Koush et al., 2015, 2013; MacInnes et al., 2016; Marins et al., 2015; McDonald et al., 2017; Megumi et al., 2015; Papoutsi et al., 2019, 2018; Scharnowski et al., 2012, 2015; Skouras & Scharnowski, 2019; Sorger, Kamp, Weiskopf, Peters, & Goebel, 2018; Spetter et al., 2017; Yao et al., 2016; Young et al., 2017).

**Figure 1:**
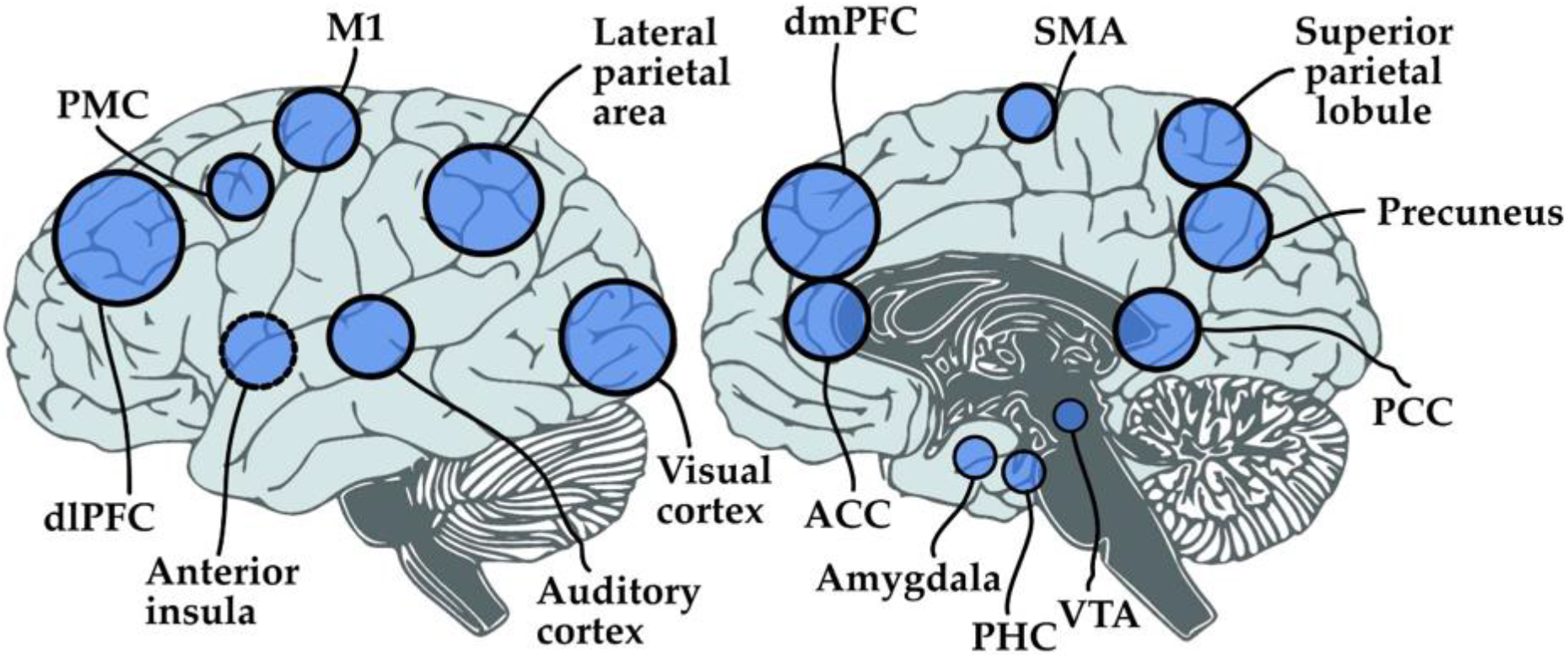
Schematic representation of areas targeted in the neurofeedback experiments. Studies included in this meta-analysis trained activity within and connectivity between more than 20 different cortical and subcortical regions of interest that are associated with various behavioral functions. This figure is for overview purposes only and does not reflect the exact coordinates or shape of the chosen ROIs. Abbreviations: ACC – Anterior Cingulate Cortex, dlPFC – dorsolateral Prefrontal Cortex, dmPFC – dorsomedial Prefrontal Cortex, M1 – Primary Motor Cortex, PCC – Posterior Cingulate Cortex, PMC – Pre-Motor Cortex, PHC – Parahippocampal Cortex, SMA – Supplementary Motor Cortex, VTA – Ventral Tegmental Area.

### Received data on pre training activity and neurofeedback learning success

We asked the authors to provide one value determining neurofeedback success for each neurofeedback training run, and one value determining pre-training brain activity levels within the ROI that was trained during neurofeedback. In particular, we asked for individual data for each participant of an experimental neurofeedback training group, excluding control groups such as receiving sham feedback or modulating brain regions of no interest. Most contributions consisted of data that were already fully analyzed and published.

For 23 studies [1-7, 9-13, 15-24] we received fully processed neurofeedback success measures for each neurofeedback training run. For reasons of simplicity and comparability to previously published results, neurofeedback success per run was defined as the measure of neurofeedback success that has been primarily assessed in the respective study and, for published data, has been previously reported in the corresponding publications. In one case (Kirschner et al. (in prep.)) [8], where raw data were provided, we calculated neurofeedback success based on standard general linear model (GLM) analyses, as described below. Overall neurofeedback learning success was then calculated based on these per-run success measures (see below).

For most studies, we also received fully processed beta values for average pre-training activity levels within the trained ROI. In some cases, we extracted these values using target ROI masks and contrast images of the corresponding pre-training run [3, 6, 9, 10, 18], or from raw data [7, 19].

### Data analysis of raw data

For the study that shared the raw data, we analyzed the data using a standard preprocessing procedure in native space (slice time correction, motion correction, co-registration, spatial smoothing with a Gaussian kernel of 6mm full width at half maximum, no normalization) using SPM12 (http://www.fil.ion.ucl.ac.uk/spm/software/spm12/). We then performed first level GLM analyses on the neurofeedback as well as the pre-training runs to model the corresponding study’s neurofeedback blocks or blocks engaging the ROI during pre-training runs, respectively.

To define pre-training activity, we extracted the average activity over all voxels within the trained ROI. When several ROIs were trained, the average over all ROIs was calculated. Activity was assessed by the beta weight representing the ROI-engaging task during pretraining. For this study, neurofeedback learning success for each neurofeedback training run was assessed in the same way, using the beta value representing the corresponding study’s neurofeedback blocks.

### Meta-analysis

To date, there is no consensus on how neurofeedback learning success should be defined and measured. Thus, in order to improve generalizability, we investigated the two most commonly used measures for assessing neurofeedback learning success (Thibault, MacPherson, Lifshitz, Roth, & Raz, 2018), namely (1) the slope of the learning curve (i.e., the regression line over the success measures for each training run), and (2) the difference between neurofeedback regulation success during the last and the first training run. Success measures of studies where participants had to perform down-regulation were multiplied by – 1. For each study, we then calculated the correlation between pre-training brain activity and these two success measures using Spearman correlations.

In addition, we investigated whether pre-training ROI activity levels might be more predictive of success during neurofeedback training runs that were performed in close temporal distance to the pre-training run. To this end, we performed correlation testing between pretraining activity levels and neurofeedback success in the very first training run.

First, we performed a meta-analysis over all the 24 studies included here. Subsequently, we repeated the meta-analysis for six different groupings of study data, to avoid confounds that may have been caused by differences between patients and healthy subjects, activity-based and connectivity-based neurofeedback paradigms, functional domains, or type of pre-training run:

1. Data from healthy subjects performing activity-based neurofeedback
2. Data from healthy subjects performing connectivity-based neurofeedback
3. Data from patients performing activity-based neurofeedback
4. Data from patients performing connectivity-based neurofeedback
5. Data split according to the functional domain of the trained ROI: i) sensory areas, ii) motor areas, iii) reward areas, iv) emotion processing area/amygdala, v) higher order cognitive processing areas/DMN and PFC)
6. Data split according to the type of pre-training run that was performed: i) functional localizer run, ii) no-feedback run, iii) ROI-engaging run that was not used for localization

Due to small sample sizes, further subdivisions of the data in (5) and (6) according to patients/healthy subjects and activity/connectivity measures were not performed. For each of these six groups as well as the entire sample (all data from all studies), we calculated overall meta-correlations using a weighted (weights based on the number of participants included in the study) random-effects model. All statistical meta-analyses were performed using the *meta* package in R using the *metacor*function (www.cran.r-project.org/web/packages/metacor). Studies that included both patients and healthy subjects, and studies that investigated both connectivity-as well as activity-based neurofeedback were split into several corresponding sub-groups accordingly. One study that trained a different ROI for each participant [21] was not considered in the ROI-based group split. Further, some of the studies included in the no-feedback group or the ROI-engaging paradigm group included a functional localizer scan in their experimental design but, due to data dropouts, the corresponding no-feedback or ROI-engaging paradigm runs were used to extract activity levels. In addition, we performed several analyses to quantify heterogeneity of effect sizes using the *Meta-Essentials* tool (Suurmond & Hak, 2017).

## Results

### Meta-analysis over all studies

The meta-analysis over the entire sample of all studies did not reveal a significant relationship between pre-training activity levels and neither of the two neurofeedback success measures (slope of the learning curve: r(27) = −0.02, p = 0.80; last vs. first run: r(27) = −0.00, p = 0.94). Further, pre-training activity levels did not show a significant correlation to neurofeedback success during the very first neurofeedback run (r(27) = 0.08, p = 0.36). Heterogeneity analyses indicated low heterogeneity of effect sizes (Higgins, Thompson, Deeks, & Altman, n.d.), both for the slope of the learning curve (Q = 30.13, Q-df = 3.13, p_Q_ = 0.31, I_2_ = 10.38%, T_2_ = 0.01, T= 0.10), and for the last vs. first neurofeedback training run (Q = 27.61, Q-df = 0.61, p_Q_ = 0.43, I_2_ = 2.20%, T_2_ = 0.00, T = 0.04). Correlations between pretraining activity levels and success in the very first neurofeedback run were moderately heterogeneous across studies (Q = 49.35, Q-df = 22.35, p_Q_ = 0.005, I_2_ = 45.29%, T_2_ = 0.07, T = 0.27). Figure 2, 3, and 4 show forest plots for correlations between pre-training activity levels and the slope success measure, the difference between the last and the first run success measures, and success during the very first neurofeedback run, respectively.

**Figure 2:**
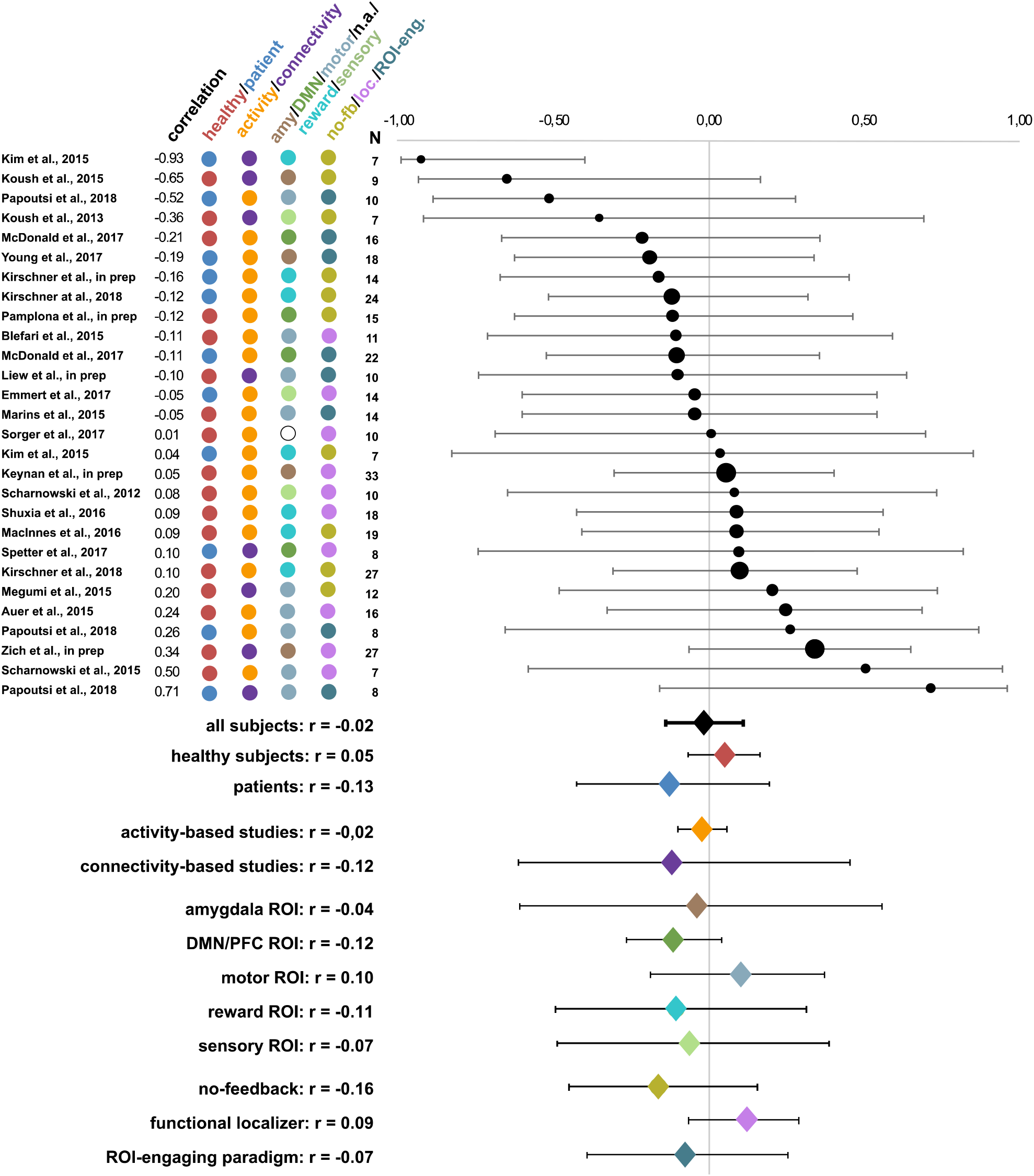
Averaged weighted Spearman correlations between pre-training activity levels and neurofeedback learning success as measured by the slope of the learning curve. Circle sizes represent the corresponding study’s sample sizes. Further, the coloring scheme reflects the corresponding grouping of the subjects (healthy subjects/patients) and the studies (type of feedback, trained target region(s) and type of pre-training activity levels). Overall, no correspondence between pre-training activity levels and neurofeedback learning success was found. Abbreviations: amy: amygdala, DMN: default mode network, n.a.: not applicable, no-fb: no feedback, loc: localizer, ROI-eng: ROI-engaging

**Figure 3:**
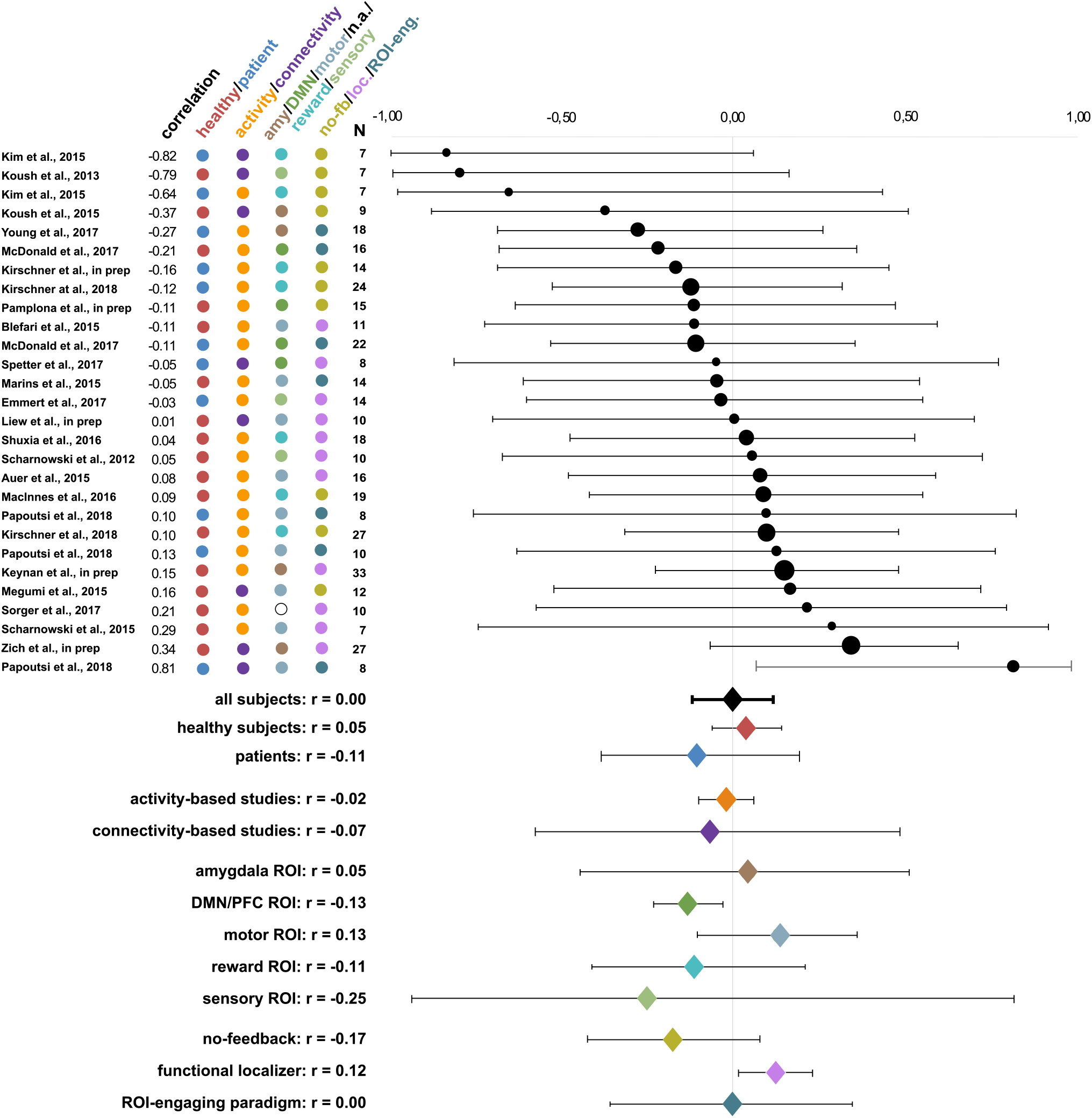
Averaged weighted Spearman correlations between pre-training activity levels and neurofeedback learning success as measured by the difference between neurofeedback success in the last and the first neurofeedback run. Circle sizes represent the corresponding study’s sample sizes. Further, the coloring scheme reflects the corresponding grouping of the subjects (healthy subjects/patients) and the studies (type of feedback, trained target region(s) and type of pre-training activity levels). Overall, no correspondence between pre-training activity levels and neurofeedback learning success was found, except for when only investigating pre-training activity levels during a functional localizer run. Abbreviations: amy: amygdala, DMN: default mode network, n.a.: not applicable, no-fb: no feedback, loc: localizer, ROI-eng: ROI-engaging

**Figure 4:**
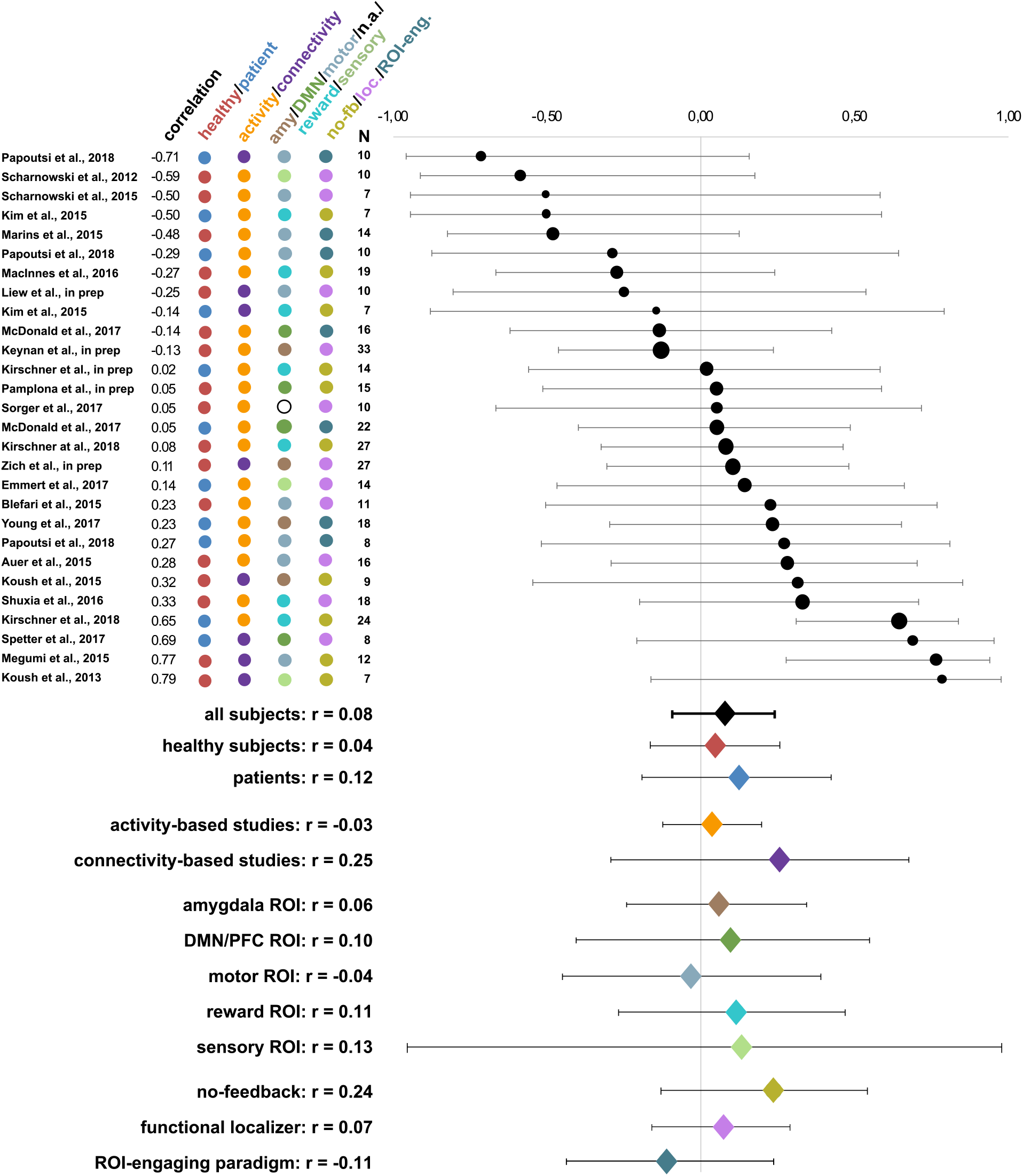
Averaged weighted Spearman correlations between pre-training activity levels and neurofeedback learning success during the first neurofeedback run. Circle sizes represent the corresponding study’s sample sizes. Further, the coloring scheme reflects the corresponding grouping of the subjects (healthy subjects/patients) and the studies (type of feedback, trained target region(s) and type of pre-training activity levels). Overall, no correspondence between pre-training activity levels and neurofeedback success in the very first neurofeedback run was found. Abbreviations: amy: amygdala, DMN: default mode network, n.a.: not applicable, no-fb: no feedback, loc: localizer, ROI-eng: ROI-engaging

### Activity-based neurofeedback with healthy subjects

For activity-based neurofeedback with healthy subjects, we found no significant relationship between pre-training activity levels and neurofeedback learning success for neither neurofeedback success measures; i.e. neither based on the slope of the regression line over all neurofeedback runs (r = 0.04, p = 0.57), nor based on the difference between the last and the first run (r = 0.04, p = 0.56). Heterogeneity measures for activity-based studies on healthy subjects were smaller than heterogeneity measures for all studies. They indicated very low heterogeneity of effect sizes, both for the slope-based (Q = 3.24, Q-df < 0, p_Q_ = 0.99, I_2_ = 0.00%, T_2_ = 0.00, T= 0.00) as well as the difference-based (Q = 2.36, Q-df < 0, p_Q_ = 1.00, I_2_ = 0.00%, T_2_ = 0.00, T = 0.00) neurofeedback learning success measure.

For activity-based neurofeedback studies with healthy subjects, we also found no significant relationship between pre-training activity levels and success in the first neurofeedback run (r = −0.06, p = 0.49), with 6 out of 12 studies even showing a negative correlation. Heterogeneity measures again showed low heterogeneity of effect sizes (Q = 12.45, Q-df < 0, p_Q_ = 0.33, I_2_ = 11.68%, T_2_ = 0.01, T= 0.10).

### Connectivity-based neurofeedback with healthy subjects

Similar to the results on activity-based neurofeedback studies with healthy subjects, for connectivity-based neurofeedback studies on healthy subjects, we again found no significant correlation between pre-training activity levels and neurofeedback learning success (slope of the regression line: r = −0.04, p = 0.85; last vs. first run difference: r = −0.06, p = 0.77). Heterogeneity measures for connectivity-based studies on healthy subjects indicated a moderate heterogeneity of effect sizes, for the slope-based (Q = 7.18, Q-df < 0, p_Q_ = 0.07, I_2_ = 58.24%, T_2_ = 0.17, T = 0.41) and for the difference-based (Q = 8.41, Q-df < 0, p_Q_ = 0.08, I_2_ = 52.43%, T_2_ = 0.13, T = 0.36) neurofeedback learning success measure.

Pre-training activity levels were slightly predictive of neurofeedback success in the very first neurofeedback run (r = 0.38, p = 0.10). Heterogeneity analyses showed again moderate heterogeneity (Q = 9.99, Q-df = 5.99, p_Q_ = 0.04, I_2_ = 59.96%, T_2_ = 0.17, T= 0.41).

### Activity-based neurofeedback with patients

For activity-based neurofeedback studies across different patient populations, we did not find a significant correlation between pre-training activity levels and neurofeedback learning success, for neither neurofeedback learning success measures (slope of the learning curve: r = −0.13, p = 0.20; last vs. first run difference: r = −0.14, p = 0.19). Here, 6 out of 8, and 7 out of 8 studies showed a slightly negative relationship, respectively. Heterogeneity of effects sizes was very low (slope: Q = 2.42, Q-df < 0, p_Q_ = 0.93, I_2_ = 0.00%, T_2_ = 0.00, T= 0.00; last vs. first difference: Q = 2.79, Q-df = 0.79, p_Q_ = 0.90, I_2_ = 0.00%, T_2_ = 0.00, T= 0.00). Pretraining activity levels in patients were also not predictive for neurofeedback success in the very first training run (r = 0.18, p = 0.18; heterogeneity measures: Q = 11.13, Q-df = 4.13, p_Q_ = 0.13, I_2_ = 37.08%, T_2_ = 0.05, T= 0.23).

### Connectivity-based neurofeedback with patients

There were only three studies of connectivity-based neurofeedback with patients. In this category, similarly to the previous ones, we observed no significant correlation between pretraining activity levels and neurofeedback learning success as defined by the slope of the learning curve (r = −0.20, p = 0.77). Heterogeneity of effect sizes, in this case, was high (Q = 14.73, Q-df = 12.73, p_Q_ = 0.00, I_2_ = 86.43%, T_2_ = 1.37, T = 1.17). Moreover, when neurofeedback learning success was defined by the difference between the last and first neurofeedback training run, we observed a significant negative correlation (r = −0.01, p = 0.984). Here, heterogeneity of effect sizes was high (Q = 11.70, Q-df = 1.70, p_Q_ = 0.003, I_2_ = 82.90%, T_2_ = 1.04, T= 1.02). Pre-training activity levels were not predictive of neurofeedback success in the very first training run (r = −0.06, p = 0.91; heterogeneity measures: Q = 7.65, Q-df = 5.65, p_Q_ = 0.02, I_2_ = 73.86%, T_2_ = 0.61, T= 0.78).

### Functional domain of the trained ROI

To investigate whether ROIs within specific functional domains would show stronger correlations than others, we clustered the studies based on the functional domain of the (main) neurofeedback traget ROI(s). For neurofeedback success, as measured by the slope of the learning curve (see Table S1 in Supporting Information), we did not find significant effects for any of the assessed functional domains, i.e. amygdala (emotion processing), DMN/PFC (mind wandering and higher cognitive functioning), motor functioning, reward processing, and other sensory domains. For neurofeedback success measured by the difference between success in the first and the last neurofeedback run (see Table S2 in Supporting Information), we found a negative correlation for studies that focused on DMN/PFC regulation (r = −0.13, p < 0.001). We did not find significant effects for any functional domain clusters when investigating the correlation between pre-training activity levels and neurofeedback success during the first neurofeedback run (see Table S3 in Supporting Information).

### Type of pre-training run

Pre-training activity levels were either based on a no-feedback run, a functional localizer run, or on another task engaging the ROI that was not used for localizing the ROI, e.g. a finger tapping task when neurofeedback training was targeting the motor cortex [13]. Overall, studies with a functional localizer run showed a significant positive correlation between the localizer activity levels and neurofeedback learning success as measured by the difference between neurofeedback learning success in the last and the first neurofeedback run (r = 0.12, p = 0.003). However, this correlation was not significant when success was measured by the slope of the learning curve (r = 0.09, p = 0.20). For activity levels during other pretraining runs we did not observe a significant correlation with learning success. Further, none of the three types of pre-training run groups showed significant correlations between pretraining activity and the very first neurofeedback run (see Table S4-S6 in Supporting Information for exact values).

## Discussion

Here, we performed a meta-analysis with 24 different fMRI-based neurofeedback studies to investigate whether pre-training activity levels can be used to predict neurofeedback learning success. In our dataset of 401 subjects undergoing neurofeedback training, we did not find an overall significant relationship between these two measures, i.e., ROI activity prior to neurofeedback training and neurofeedback learning success were not significantly correlated.

One of the reasons for not having found an overall relationship between pre-training activity and learning success might be that the studies included in this meta-analysis are quite diverse in terms of, for example, the research question, the target ROI, the feedback signal and the population. One the other hand, heterogeneity analyses of effect sizes across all studies revealed that our sample of studies was sufficiently homogenous for a meta-analysis and that the result was unlikely to be confounded by single studies. Nevertheless, we aimed at partly mitigating heterogeneity by repeating the analysis for different groups containing only healthy participants or patients, activity- or connectivity-based neurofeedback, only studies with the same type of pre-training run, and by grouping studies based on the functional domain of the trained ROI. Unfortunately, further subclassifications in, for instance, studies who performed up-vs. down-regulation could not be performed due to too low sample sizes.

### Differences between healthy subjects and patients

Neither healthy subjects nor patients showed a significant correlation between pre-training activity levels and neurofeedback learning success.

Interestingly, the majority of patient studies showed a negative correlation between neurofeedback learning success and pre-training activity levels, while we observed more positive correlations for studies with healthy subjects. This might be explained by the fact that some disorders of the patient populations included in this meta-analysis are associated with hyper-activation in the targeted ROIs, thus making it more difficult to increase ROI activity even further. For instance, patients with schizophrenia show a hyper-activation in the VTA (Howes & Kapur, 2009) and might therefore experience ceiling effects when trying to upregulate their VTA activity. Likewise, symptom severity might be associated with increased ROI activity, which again can influence a patient’s neurofeedback learning performance. For example, patients suffering from substance use disorder who show highly increased craving-induced brain activity levels might be less successful in down-regulating craving-related brain signals than addiction patients who only show mildly increased craving-related brain activity. Increased brain activity levels in higher order brain areas might also be an indicator for decreased cognitive capacities – as the performed task constitutes a particular challenge to the patients, they might experience exhaustion during the following neurofeedback training runs. Further, aspects like differences in adaptation, motivation, deficits in sustained attention etc. that are often reported in specific patient populations, might also drive neurofeedback training success differences.

### Activity-vs. connectivity-based neurofeedback

Neither activity-nor connectivity-based neurofeedback studies showed a significant correlation between pre-training activity levels and neurofeedback learning success. Moreover, while heterogeneity measures for effect sizes of activity-based neurofeedback studies showed very low heterogeneity, this was not the case for effect sizes of connectivity-based neurofeedback studies. Here, heterogeneity measures of effect sizes revealed moderate heterogeneity, indicating that effect sizes in connectivity-based studies might be too diverse to be grouped together in one meta-analysis. This might be related to connectivity-based neurofeedback studies still being sparse with overall limited sample sizes. Another confounding factor might be that for connectivity-based studies pre-training activity levels are not as similar to neurofeedback success measures as for activity-based studies. Consequently, future studies should investigate whether pre-training levels based on connectivity are more predictive for neurofeedback learning success in connectivity-based neurofeedback studies and, in addition, whether effect sizes based on pre-training connectivity levels are less heterogeneous._In fact, a recent study <u>found that DMN upregulation learning and down-regulation learning scores are partly determined by pre-neurofeedback resting-state eigenvector centrality of the PCC/PCu</u> (Skouras & Scharnowski, 2019). Further, another study observed resting state connectivity to be predictive for neurofeedback learning success in patients with obsessive-compulsive disorder (Scheinost et al., 2014). These findings should be replicated and tested to assess whether they are generalizable.

### Type of pre-training run

Interestingly, when grouping together studies based on the paradigm of the run during which pre-training activity levels were collected, we observed a significant positive correlation between pre-training activity levels and neurofeedback success (as measured by the difference in neurofeedback success between the last and the first neurofeedback run) for studies with a functional localizer run. This indicates that pre-training activity levels might indeed be linked to neurofeedback learning success, but only when the neurofeedback target ROI is completely activated during the pre-training run, as it is the case in functional localizer runs. In contrast, in no-feedback and other ROI-engaging paradigms (i.e. not functional localizers), the target ROI may be engaged by the pre-training paradigm, however some voxels within the specified ROI may not be specifically involved in the neural processes under investigation. Consequently, when extracting pre-training activity levels from no-feedback and ROI-engaging (but not functional localizer) pre-training runs, more voxels than those that reliably activate during the performed task contribute to the derived signal and, thus, limit its predictive power for neurofeedback success.

In contrast to functional localizer runs, no-feedback runs (i.e. where the participants were performing the same task as during a neurofeedback run but without getting any feedback) did not predict neurofeedback learning success. Surprisingly, the no-feedback runs performed just before the neurofeedback training commenced were not even predictive of performance during the very first neurofeedback training run. No-feedback runs (also referred to as “transfer runs” when performed after neurofeedback training) are identical to the neurofeedback training runs (i.e. same ROI, same design, similar instructions, similar mental task, etc.) except that no feedback signal is presented. This indicates that providing feedback might only be a small experimental addition, but one that changes the paradigm significantly. Previous studies have already highlighted the discrepancy between pre-training no-feedback runs and neurofeedback runs by showing that no-feedback runs differ substantially from neurofeedback training runs in terms of functional connectivity changes (Haller et al., 2013), self-regulation performance (Robineau et al., 2014), and signal-to-noise ratio (Papageorgiou, Lisinski, McHenry, White, & LaConte, 2013). This also indicates that the feedback has a stronger effect on neurofeedback training success than the actual task the participant is performing in the scanner. Indeed, recent implicit neurofeedback studies show that neurofeedback learning is possible even when participants are not informed what the neurofeedback signal represents and are not provided with mental strategy instructions that are related to the function of the target ROI (Cortese, Amano, Koizumi, Kawato, & Lau, 2016; Koizumi et al., 2017; Shibata et al., 2011; Taschereau-Dumouchel et al., 2018). These findings show that neurofeedback runs are special in that they constitute their own specific experimental condition that is distinct from seemingly-related conditions such as transfer runs without neurofeedback. Thus, they should be analyzed separately and, for example, performance during no-feedback and training runs should not be combined in one continuous learning curve.

### Neurofeedback learning measure

For the purpose of generalizability, we assessed neurofeedback learning success by (a) the slope of the regression line over the per-run success measures, and (b) the difference between neurofeedback success during the last run compared to the first neurofeedback run. These two measures are frequently used in neurofeedback studies and they capture the efficiency of neurofeedback learning (slope) as well as the effect of neurofeedback learning (difference between the last and first run). These two measures are highly correlated with an average correlation of r = 0.78 across all studies. However, in the neurofeedback literature there is still no generally accepted best measure for assessing neurofeedback learning success. Other potential success measures are, for example, the difference between pre- and post-training transfer runs (e.g. Auer et al., 2015; Koush et al., 2015; MacInnes et al., 2016; Megumi et al., 2015), or the behavioral/clinical improvements (e.g. deCharms et al., 2005; Emmert, Kopel, Koush, Maire, Senn, De Ville, et al., 2017; Linden et al., 2012; Scharnowski et al., 2015; Young et al., 2017). One might speculate that predictions might have been better if we had used an alternative neurofeedback learning measure. On the other hand, pre-training activity was not even predictive of the very first neurofeedback training run activity (Supporting Information Table S6) and for this comparison identical measures (i.e. ROI activity) were used.

The underlying problem with respect to defining a commonly accepted neurofeedback learning measure is that there is no established model of neurofeedback learning (Sitaram et al., 2016), thus making it difficult to define the key parameters involved in successful neurofeedback training. Also, individual learning curves are quite diverse so that defining a one-fits-all learning measures that captures the multitude of manifestations of neurofeedback learning is very challenging. For that reason, running the analyses in parallel for two different neurofeedback performance measures is a pragmatic solution aiming to capture potential predictors of learning success.

### Predicting neurofeedback learning success

Overall, we were not able to predict neurofeedback learning success from pre-training activity levels. However, when observing only studies that defined their neurofeedback target ROI(s) based on a functional localizer task, we identified a positive correlation between pretraining activity levels and neurofeedback learning success (i.e. slope-based and difference-based). These results indicate that neurofeedback performance is connected to pre-training activity levels, but only when all neurofeedback target voxels can be actively engaged by the functional pre-training task. Nevertheless, even for this group of neurofeedback training studies, we did not find any significant results for individual studies. Further, the weak correlation of r = 0.12 indicates that it is not possible to create considerably accurate predictions on which participants might be able to perform well during neurofeedback training and which participants will fail to do so.

Taken together, factors that can already be assessed in pre-training transfer and localizer runs, such as noise levels, participant compliance, or the responsiveness of a particular ROI, are not the main causes for the large inter-individual differences in neurofeedback learning success (Bray et al., 2007; Chiew et al., 2012; deCharms et al., 2005; Johnson et al., 2012; Robineau et al., 2014; Scharnowski et al., 2012; Yoo et al., 2008).

This poses the question of what other information might be useful as a predictor for neurofeedback learning success. Obvious candidates would be standardized questionnaires or behavioral tasks that could be used for participant selection without having to acquire imaging data. Unfortunately, evidence for the predictive power of such measures is sparse. While two studies found that the pain Coping Strategies Questionnaire (Rosenstiel & Keefe, 1983) and state anxiety scores (Spielberger, 2010) predict success in learning to regulate the ACC (Emmert, Breimhorst, et al., 2017) and emotion networks (Koush et al., 2015), respectively, another study did not find correlations between pre-training spatial orientation (Stumpf & Fay, 1983), creative imagination (Barber & Wilson, 1978), or mood scores (Zerssen, 1976) scores and success in learning to regulate pre-motor and parahippocampal ROIs (Scharnowski et al., 2015). A recent systematic review on psychological factors that might influence neurofeedback learning success in EEG and fMRI studies argues that factors such as attention and motivation might play an important role in successful neurofeedback training runs (Cohen & Staunton, 2019). However, although these are likely candidates for affecting neurofeedback learning, a concrete empirical effect of these factors has so far only been reported in one fMRI-based neurofeedback study (Chiew et al., 2012), showing a clear necessity for more empirical investigations on these factors.

In EEG neurofeedback, several factors have been observed to be correlated with neurofeedback learning success (Alkoby et al., 2017), but they were only reported in single EEG studies have not yet been tested in fMRI-based neurofeedback studies. For instance, factors that seemed to have a positive influence on EEG-based neurofeedback learning success were regular spiritual practice (Kober et al., 2017) or a relaxing attitude towards one’s ability to control technological devices (Witte, Kober, Ninaus, Neuper, & Wood, 2013). Also, other brain-based measures that are, for example, focused on areas more generally involved in self-regulation (Emmert et al., 2016) or on connectivity rather than activity levels might be suitable candidates that should be explored in future studies. The latter might be particularly relevant for connectivity-based neurofeedback studies, but we were not able to test this due to lack of suitable data. Further possible candidates for predicting neurofeedback success might be factors that have already been identified to be predictive of cognitive and behavioral training success in non-neurofeedback studies, e.g. activity in areas related to stimulus encoding and motor control has been found to be predictive of motor learning (Herholz, Coffey, Pantev, & Zatorre, 2016), and activity in the motor network has been found to predict training-related changes in working memory (Simmonite & Polk, 2019). Finally, very recent work by Skouras et al. indicates that neurofeedback learning performance can be influenced by biological factors such as genetic and anatomical predispositions (Skouras et al., 2019), thus demonstrating the complexity of the underlying processes and the need for using multi-modal datasets.

Hence, currently, no robust predictors for neurofeedback learning success have been identified, and, even if predictions can be made, they are likely study-specific (i.e. questionnaires that are specific to the trained ROI) and might not generalize across studies.

## Conclusion

Here, we aimed at finding general pre-training predictors for neurofeedback training success. We observed a slightly positive correlation between pre-training activity levels during a functional localizer run and neurofeedback learning success, but we were not able to identify common brain-based success predictors across our diverse cohort of studies. In order to achieve the goal of finding predictors for neurofeedback learning success advances need to be made: in developing (1) models for neurofeedback learning, (2) establishing robust measures for neurofeedback learning, and (3) in increasing the database including acquired candidate measures across numerous studies. The reward of such a joint effort would be increased efficacy and cost-effectiveness of this promising scientific and therapeutic method.

## Acknowledgements

This work was supported by the Forschungskredit of the University of Zurich, the Foundation for Research in Science and the Humanities at the University of Zurich (STWF-17-012), the Baugarten Stiftung, and the Swiss National Science Foundation (BSSG10_155915, 32003B_166566). The authors declare no conflicts of interest.

