## Supporting Information for "Can we predict real-time fMRI neurofeedback learning success from pre-training brain activity?"

| **Study name / Subgroup name** | **Correlation** | **CI Lower limit** | **CI Upper limit** | **Weight** | **Q** | **p_Q_** | **I^2^** | **T^2^** | **T** |
| --- | --- | --- | --- | --- | --- | --- | --- | --- | --- |
| Keynan et al., in prep | 0.05 | -0.31 | 0.40 | 31.58% |  |  |  |  |  |
| Koush_et al., 2015 | -0.65 | -0.94 | 0.16 | 14.55% |  |  |  |  |  |
| Young et al., 2017 | -0.19 | -0.63 | 0.34 | 24.43% |  |  |  |  |  |
| Zich et al., in prep | 0.34 | -0.07 | 0.65 | 29.43% |  |  |  |  |  |
| **amygdala** | **-0.04** | **-0.61** | **0.56** | **3.71%** | **7.18** | **0.07** | **0.58** | **0.08** | **0.28** |
| McDonald et al., 2017 | -0.21 | -0.67 | 0.36 | 26.53% |  |  |  |  |  |
| McDonald et al., 2017 | -0.11 | -0.52 | 0.35 | 38.78% |  |  |  |  |  |
| Pamplona et al., in prep | -0.12 | -0.63 | 0.46 | 24.49% |  |  |  |  |  |
| Spetter et al., 2017 | 0.10 | -0.75 | 0.82 | 10.20% |  |  |  |  |  |
| **DMN/PFC** | **-0.12** | **-0.27** | **0.04** | **67.01%** | **0.36** | **0.95** | **0.00** | **0.00** | **0.00** |
| Auer et al., 2015 | 0.24 | -0.33 | 0.69 | 18.17% |  |  |  |  |  |
| Blefari et al., 2015 | -0.11 | -0.71 | 0.59 | 11.65% |  |  |  |  |  |
| Megumi et al., 2015 | 0.20 | -0.48 | 0.73 | 13.00% |  |  |  |  |  |
| Liew et al., in prep | -0.10 | -0.74 | 0.64 | 10.28% |  |  |  |  |  |
| Marins et al., 2015 | -0.05 | -0.60 | 0.54 | 15.63% |  |  |  |  |  |
| Papoutsi et al., 2018 | -0.52 | -0.89 | 0.28 | 10.28% |  |  |  |  |  |
| Papoutsi et al., 2018 | 0.26 | -0.66 | 0.87 | 7.47% |  |  |  |  |  |
| Papoutsi et al., 2018 | 0.71 | -0.16 | 0.96 | 7.47% |  |  |  |  |  |
| Scharnowski et al., 2015 | 0.50 | -0.58 | 0.94 | 6.03% |  |  |  |  |  |
| **Motor ROI** | **0.10** | **-0.18** | **0.37** | **10.42%** | **8.54** | **0.38** | **0.06** | **0.01** | **0.09** |
| Kim et al., 2015 | 0.04 | -0.83 | 0.85 | 7.04% |  |  |  |  |  |
| Kim et al., 2015 | -0.93 | -0.99 | -0.40 | 7.04% |  |  |  |  |  |
| Kirschner et al., 2018 | -0.12 | -0.52 | 0.32 | 18.92% |  |  |  |  |  |
| Kirschner et al., 2018 | 0.10 | -0.31 | 0.48 | 19.91% |  |  |  |  |  |
| Kirschner et al., in prep | -0.16 | -0.67 | 0.45 | 13.91% |  |  |  |  |  |
| MacInnes et al., 2016 | 0.09 | -0.41 | 0.55 | 16.84% |  |  |  |  |  |
| Shuxia et al., 2016 | 0.09 | -0.43 | 0.56 | 16.33% |  |  |  |  |  |
| **Reward ROI** | **-0.11** | **-0.50** | **0.31** | **5.19%** | **11.62** | **0.07** | **0.48** | **0.07** | **0.27** |
| Emmert et al., 2017 | -0.05 | -0.60 | 0.54 | 50.00% |  |  |  |  |  |
| Koush et al., 2013 | -0.36 | -0.92 | 0.69 | 18.18% |  |  |  |  |  |
| Scharnowski et al., 2012 | 0.08 | -0.65 | 0.73 | 31.82% |  |  |  |  |  |
| **Sensory ROI** | **-0.07** | **-0.49** | **0.39** | **13.66%** | **0.53** | **0.77** | **0.00** | **0.00** | **0.00** |

Table S1: **Averaged weighted Spearman correlations between pre-training activity levels and neurofeedback learning success as measured by the slope of the learning curve, clustered by the functional domain of the trained ROI or the main ROI of the trained ROIs** Abbreviations: DMN – default mode network, PFC: prefrontal cortex

| **Study name / Subgroup name** | **Correlation** | **CI Lower limit** | **CI Upper limit** | **Weight** | **Q** | **p_Q_** | **I^2^** | **T^2^** | **T** |
| --- | --- | --- | --- | --- | --- | --- | --- | --- | --- |
| Keynan et al., in prep | 0.15 | -0.22 | 0.48 | 33.81% |  |  |  |  |  |
| Koush_et al., 2015 | -0.37 | -0.87 | 0.51 | 12.25% |  |  |  |  |  |
| Young et al., 2017 | -0.27 | -0.68 | 0.26 | 23.48% |  |  |  |  |  |
| Zich et al., in prep | 0.34 | -0.06 | 0.65 | 30.46% |  |  |  |  |  |
| **amygdala** | **0.05** | **-0.44** | **0.51** | **3.21%** | **5.20** | **0.16** | **0.42** | **0.04** | **0.21** |
| McDonald et al., 2017 | -0.21 | -0.67 | 0.36 | 26.53% |  |  |  |  |  |
| McDonald et al., 2017 | -0.11 | -0.52 | 0.35 | 38.78% |  |  |  |  |  |
| Pamplona et al., in prep | -0.11 | -0.62 | 0.47 | 24.49% |  |  |  |  |  |
| Spetter et al., 2017 | -0.05 | -0.80 | 0.77 | 10.20% |  |  |  |  |  |
| **DMN/PFC** | **-0.13** | **-0.23** | **-0.03** | **84.49%** | **0.15** | **0.99** | **0.00** | **0.00** | **0.00** |
| Auer et al., 2015 | 0.08 | -0.47 | 0.59 | 18.84% |  |  |  |  |  |
| Blefari et al., 2015 | -0.11 | -0.71 | 0.59 | 11.59% |  |  |  |  |  |
| Megumi et al., 2015 | 0.16 | -0.51 | 0.72 | 13.04% |  |  |  |  |  |
| Liew et al., in prep | 0.01 | -0.69 | 0.70 | 10.14% |  |  |  |  |  |
| Marins et al., 2015 | -0.05 | -0.60 | 0.54 | 15.94% |  |  |  |  |  |
| Papoutsi et al., 2018 | 0.13 | -0.62 | 0.75 | 10.14% |  |  |  |  |  |
| Papoutsi et al., 2018 | 0.10 | -0.75 | 0.82 | 7.25% |  |  |  |  |  |
| Papoutsi et al., 2018 | -0.81 | -0.97 | -0.07 | 7.25% |  |  |  |  |  |
| Scharnowski et al., 2015 | 0.29 | -0.73 | 0.91 | 5.80% |  |  |  |  |  |
| **Motor ROI** | **0.13** | **-0.10** | **0.36** | **6.56%** | **6.02** | **0.65** | **0.00** | **0.00** | **0.00** |
| Kim et al., 2015 | -0.64 | -0.96 | 0.43 | 5.51% |  |  |  |  |  |
| Kim et al., 2015 | -0.82 | -0.98 | 0.06 | 5.51% |  |  |  |  |  |
| Kirschner et al., 2018 | -0.12 | -0.52 | 0.32 | 20.45% |  |  |  |  |  |
| Kirschner et al., 2018 | 0.10 | -0.31 | 0.48 | 22.22% |  |  |  |  |  |
| Kirschner et al., in prep | -0.16 | -0.67 | 0.45 | 12.95% |  |  |  |  |  |
| MacInnes et al., 2016 | 0.09 | -0.41 | 0.55 | 17.05% |  |  |  |  |  |
| Shuxia et al., 2016 | 0.04 | -0.47 | 0.53 | 16.30% |  |  |  |  |  |
| **Reward ROI** | **-0.11** | **-0.41** | **0.21** | **4.92%** | **8.10** | **0.23** | **0.26** | **0.03** | **0.16** |
| Emmert et al., 2017 | -0.03 | -0.59 | 0.55 | 41.93% |  |  |  |  |  |
| Koush et al., 2013 | -0.79 | -0.98 | 0.16 | 24.23% |  |  |  |  |  |
| Scharnowski et al., 2012 | 0.05 | -0.66 | 0.72 | 33.85% |  |  |  |  |  |
| **Sensory ROI** | **-0.25** | **-0.93** | **0.82** | **0.82%** | **3.72** | **0.16** | **0.46** | **0.13** | **0.36** |

Table S2: **Averaged weighted Spearman correlations between pre-training activity levels and neurofeedback learning success as measured by the difference between neurofeedback success in the last and the first neurofeedback run, clustered by the functional domain of the trained ROI or the main ROI of the trained ROIs** Abbreviations: DMN – default mode network, PFC: prefrontal cortex

| **Study name / Subgroup name** | **Correlation** | **CI Lower limit** | **CI Upper limit** | **Weight** | **Q** | **p_Q_** | **I^2^** | **T^2^** | **T** |
| --- | --- | --- | --- | --- | --- | --- | --- | --- | --- |
| Keynan et al., in prep | -0.13 | -0.46 | 0.24 | 40.00% |  |  |  |  |  |
| Koush_et al., 2015 | 0.32 | -0.55 | 0.85 | 8.00% |  |  |  |  |  |
| Young et al., 2017 | 0.23 | -0.30 | 0.65 | 20.00% |  |  |  |  |  |
| Zich et al., in prep | 0.11 | -0.30 | 0.48 | 32.00% |  |  |  |  |  |
| **amygdala** | **0.06** | **-0.24** | **0.34** | **50.96%** | **2.04** | **0.56** | **0.00** | **0.00** | **0.00** |
| McDonald et al., 2017 | -0.14 | -0.62 | 0.43 | 26.89% |  |  |  |  |  |
| McDonald et al., 2017 | 0.05 | -0.40 | 0.49 | 36.31% |  |  |  |  |  |
| Pamplona et al., in prep | 0.05 | -0.51 | 0.59 | 25.17% |  |  |  |  |  |
| Spetter et al., 2017 | 0.69 | -0.21 | 0.96 | 11.62% |  |  |  |  |  |
| **DMN/PFC** | **0.10** | **-0.40** | **0.55** | **17.00%** | **3.58** | **0.31** | **0.16** | **0.02** | **0.13** |
| Auer et al., 2015 | 0.28 | -0.29 | 0.71 | 13.62% |  |  |  |  |  |
| Blefari et al., 2015 | 0.23 | -0.51 | 0.77 | 11.65% |  |  |  |  |  |
| Megumi et al., 2015 | 0.77 | 0.28 | 0.94 | 12.16% |  |  |  |  |  |
| Liew et al., in prep | -0.25 | -0.80 | 0.54 | 11.06% |  |  |  |  |  |
| Marins et al., 2015 | -0.48 | -0.83 | 0.13 | 12.98% |  |  |  |  |  |
| Papoutsi et al., 2018 | 0.27 | -0.52 | 0.81 | 11.06% |  |  |  |  |  |
| Papoutsi et al., 2018 | -0.29 | -0.87 | 0.64 | 9.51% |  |  |  |  |  |
| Papoutsi et al., 2018 | -0.71 | -0.96 | 0.16 | 9.51% |  |  |  |  |  |
| Scharnowski et al., 2015 | -0.50 | -0.94 | 0.58 | 8.47% |  |  |  |  |  |
| **Motor ROI** | **-0.04** | **-0.45** | **0.39** | **12.12%** | **20.55** | **0.01** | **0.61** | **0.21** | **0.46** |
| Kim et al., 2015 | -0.50 | -0.94 | 0.59 | 7.85% |  |  |  |  |  |
| Kim et al., 2015 | -0.14 | -0.88 | 0.79 | 7.85% |  |  |  |  |  |
| Kirschner et al., 2018 | 0.65 | 0.31 | 0.84 | 18.25% |  |  |  |  |  |
| Kirschner et al., 2018 | 0.08 | -0.32 | 0.46 | 18.98% |  |  |  |  |  |
| Kirschner et al., in prep | 0.02 | -0.56 | 0.59 | 14.22% |  |  |  |  |  |
| MacInnes et al., 2016 | -0.27 | -0.67 | 0.24 | 16.63% |  |  |  |  |  |
| Shuxia et al., 2016 | 0.33 | -0.20 | 0.71 | 16.22% |  |  |  |  |  |
| **Reward ROI** | **0.11** | **-0.27** | **0.47** | **17.94%** | **14.17** | **0.03** | **0.58** | **0.11** | **0.32** |
| Emmert et al., 2017 | 0.14 | -0.47 | 0.66 | 37.36% |  |  |  |  |  |
| Koush et al., 2013 | 0.79 | -0.16 | 0.98 | 28.65% |  |  |  |  |  |
| Scharnowski et al., 2012 | -0.59 | -0.91 | 0.18 | 33.99% |  |  |  |  |  |
| **Sensory ROI** | **0.13** | **-0.96** | **0.98** | **1.97%** | **7.85** | **0.02** | **0.75** | **0.43** | **0.66** |

Table S3: **Averaged weighted Spearman correlations between pre-training activity levels and neurofeedback learning success as measured by neurofeedback success in the first neurofeedback run, clustered by the functional domain of the trained ROI or the main ROI of the trained ROIs** Abbreviations: DMN – default mode network, PFC: prefrontal cortex

| **Study name / Subgroup name** | **Correlation** | **CI Lower limit** | **CI Upper limit** | **Weight** | **Q** | **p_Q_** | **I^2^** | **T^2^** | **T** |
| --- | --- | --- | --- | --- | --- | --- | --- | --- | --- |
| Auer et al., 2015 | 0.24 | -0.33 | 0.69 | 9.42% |  |  |  |  |  |
| Blefari et al., 2015 | -0.11 | -0.71 | 0.59 | 5.80% |  |  |  |  |  |
| Emmert et al., 2017 | -0.05 | -0.60 | 0.54 | 7.97% |  |  |  |  |  |
| Keynan et al., in prep | 0.05 | -0.31 | 0.40 | 21.74% |  |  |  |  |  |
| Liew et al., in prep | -0.10 | -0.74 | 0.64 | 5.07% |  |  |  |  |  |
| Papoutsi et al., 2018 | -0.52 | -0.89 | 0.28 | 5.07% |  |  |  |  |  |
| Scharnowski et al., 2015 | 0.50 | -0.58 | 0.94 | 2.90% |  |  |  |  |  |
| Scharnowski et al., 2012 | 0.08 | -0.65 | 0.73 | 5.07% |  |  |  |  |  |
| Shuxia et al., 2016 | 0.09 | -0.43 | 0.56 | 10.87% |  |  |  |  |  |
| Sorger et al., 2017 | 0.01 | -0.69 | 0.70 | 5.07% |  |  |  |  |  |
| Spetter et al., 2017 | 0.10 | -0.75 | 0.82 | 3.62% |  |  |  |  |  |
| Zich et al., in prep | 0.34 | -0.07 | 0.65 | 17.39% |  |  |  |  |  |
| **Functional localizer** | **0.09** | **-0.06** | **0.23** | **56.43%** | **6.79** | **0.82** | **0.00** | **0.00** | **0.00** |
| Megumi et al., 2015 | 0.20 | -0.48 | 0.73 | 9.65% |  |  |  |  |  |
| Kim et al., 2015 | 0.04 | -0.83 | 0.85 | 5.38% |  |  |  |  |  |
| Kim et al., 2015 | -0.93 | -0.99 | -0.40 | 5.38% |  |  |  |  |  |
| Kirschner et al., 2018 | -0.12 | -0.52 | 0.32 | 15.15% |  |  |  |  |  |
| Kirschner et al., 2018 | 0.10 | -0.31 | 0.48 | 16.01% |  |  |  |  |  |
| Kirschner et al., in prep | -0.16 | -0.67 | 0.45 | 10.91% |  |  |  |  |  |
| Koush et al., 2013 | -0.36 | -0.92 | 0.69 | 5.38% |  |  |  |  |  |
| Koush_et al., 2015 | -0.65 | -0.94 | 0.16 | 7.32% |  |  |  |  |  |
| MacInnes et al., 2016 | 0.09 | -0.41 | 0.55 | 13.36% |  |  |  |  |  |
| Pamplona et al., in prep | -0.12 | -0.63 | 0.46 | 11.47% |  |  |  |  |  |
| **No-feedback run** | **-0.16** | **-0.45** | **0.16** | **20.77%** | **15.09** | **0.09** | **0.40** | **0.06** | **0.25** |
| Marins et al., 2015 | -0.05 | -0.60 | 0.54 | 16.61% |  |  |  |  |  |
| McDonald et al., 2017 | -0.21 | -0.67 | 0.36 | 19.20% |  |  |  |  |  |
| McDonald et al., 2017 | -0.11 | -0.52 | 0.35 | 26.32% |  |  |  |  |  |
| Papoutsi et al., 2020 | 0.26 | -0.66 | 0.87 | 8.10% |  |  |  |  |  |
| Papoutsi et al., 2020 | 0.71 | -0.16 | 0.96 | 8.10% |  |  |  |  |  |
| Young et al., 2017 | -0.19 | -0.63 | 0.34 | 21.67% |  |  |  |  |  |
| **ROI-engaging run** | **-0.03** | **-0.36** | **0.31** | **22.80%** | **5.70** | **0.34** | **0.12** | **0.01** | **0.11** |

Table S4: **Averaged weighted Spearman correlations between pre-training activity levels and neurofeedback learning success as measured by the slope of the learning curve, clustered by the paradigm of the pre-training run**

| **Study name / Subgroup name** | **Correlation** | **CI Lower limit** | **CI Upper limit** | **Weight** | **Q** | **p_Q_** | **I^2^** | **T^2^** | **T** |
| --- | --- | --- | --- | --- | --- | --- | --- | --- | --- |
| Auer et al., 2015 | 0.08 | -0.47 | 0.59 | 9.42% |  |  |  |  |  |
| Blefari et al., 2015 | -0.11 | -0.71 | 0.59 | 5.80% |  |  |  |  |  |
| Emmert et al., 2017 | -0.03 | -0.59 | 0.55 | 7.97% |  |  |  |  |  |
| Keynan et al., in prep | 0.15 | -0.22 | 0.48 | 21.74% |  |  |  |  |  |
| Liew et al., in prep | 0.01 | -0.69 | 0.70 | 5.07% |  |  |  |  |  |
| Papoutsi et al., 2018 | 0.13 | -0.62 | 0.75 | 5.07% |  |  |  |  |  |
| Scharnowski et al., 2015 | 0.29 | -0.73 | 0.91 | 2.90% |  |  |  |  |  |
| Scharnowski et al., 2012 | 0.05 | -0.66 | 0.72 | 5.07% |  |  |  |  |  |
| Shuxia et al., 2016 | 0.04 | -0.47 | 0.53 | 10.87% |  |  |  |  |  |
| Sorger et al., 2017 | 0.21 | -0.56 | 0.79 | 5.07% |  |  |  |  |  |
| Spetter et al., 2017 | -0.05 | -0.80 | 0.77 | 3.62% |  |  |  |  |  |
| Zich et al., in prep | 0.34 | -0.06 | 0.65 | 17.39% |  |  |  |  |  |
| **Functional localizer** | **0.12** | **0.03** | **0.21** | **48.77%** | **2.59** | **1.00** | **0.00** | **0.00** | **0.00** |
| Megumi et al., 2015 | 0.16 | -0.51 | 0.72 | 9.27% |  |  |  |  |  |
| Kim et al., 2015 | -0.64 | -0.96 | 0.43 | 4.78% |  |  |  |  |  |
| Kim et al., 2015 | -0.82 | -0.98 | 0.06 | 4.78% |  |  |  |  |  |
| Kirschner et al., 2018 | -0.12 | -0.52 | 0.32 | 16.23% |  |  |  |  |  |
| Kirschner et al., 2018 | 0.10 | -0.31 | 0.48 | 17.46% |  |  |  |  |  |
| Kirschner et al., in prep | -0.16 | -0.67 | 0.45 | 10.74% |  |  |  |  |  |
| Koush et al., 2013 | -0.79 | -0.98 | 0.16 | 4.78% |  |  |  |  |  |
| Koush_et al., 2015 | -0.37 | -0.87 | 0.51 | 6.74% |  |  |  |  |  |
| MacInnes et al., 2016 | 0.09 | -0.41 | 0.55 | 13.80% |  |  |  |  |  |
| Pamplona et al., in prep | -0.11 | -0.62 | 0.47 | 11.41% |  |  |  |  |  |
| **No-feedback run** | **-0.17** | **-0.43** | **0.11** | **30.25%** | **12.53** | **0.18** | **0.28** | **0.04** | **0.19** |
| Marins et al., 2015 | -0.05 | -0.60 | 0.54 | 17.26% |  |  |  |  |  |
| McDonald et al., 2017 | -0.21 | -0.67 | 0.36 | 19.02% |  |  |  |  |  |
| McDonald et al., 2017 | -0.11 | -0.52 | 0.35 | 23.14% |  |  |  |  |  |
| Papoutsi et al., 2020 | 0.10 | -0.75 | 0.82 | 10.00% |  |  |  |  |  |
| Papoutsi et al., 2020 | 0.81 | 0.07 | 0.97 | 10.00% |  |  |  |  |  |
| Young et al., 2017 | -0.27 | -0.68 | 0.26 | 20.57% |  |  |  |  |  |
| **ROI-engaging run** | **-0.01** | **-0.43** | **0.42** | **20.98%** | **8.24** | **0.14** | **0.39** | **0.06** | **0.24** |

Table S5: **Averaged weighted Spearman correlations between pre-training activity levels and neurofeedback learning success as measured by the difference in neurofeedback learning success between the last and the first neurofeedback run, clustered by the paradigm of the pre-training run**

| **Study name / Subgroup name** | **Correlation** | **CI Lower limit** | **CI Upper limit** | **Weight** | **Q** | **p_Q_** | **I^2^** | **T^2^** | **T** |
| --- | --- | --- | --- | --- | --- | --- | --- | --- | --- |
| Auer et al., 2015 | 0.28 | -0.29 | 0.71 | 9.72% |  |  |  |  |  |
| Blefari et al., 2015 | 0.23 | -0.51 | 0.77 | 6.38% |  |  |  |  |  |
| Emmert et al., 2017 | 0.14 | -0.47 | 0.66 | 8.44% |  |  |  |  |  |
| Keynan et al., in prep | -0.13 | -0.46 | 0.24 | 18.57% |  |  |  |  |  |
| Liew et al., in prep | -0.25 | -0.80 | 0.54 | 5.65% |  |  |  |  |  |
| Papoutsi et al., 2018 | 0.27 | -0.52 | 0.81 | 5.65% |  |  |  |  |  |
| Scharnowski et al., 2015 | -0.50 | -0.94 | 0.58 | 3.36% |  |  |  |  |  |
| Scharnowski et al., 2012 | -0.59 | -0.91 | 0.18 | 5.65% |  |  |  |  |  |
| Shuxia et al., 2016 | 0.33 | -0.20 | 0.71 | 10.95% |  |  |  |  |  |
| Sorger et al., 2017 | 0.05 | -0.66 | 0.72 | 5.65% |  |  |  |  |  |
| Spetter et al., 2017 | 0.69 | -0.21 | 0.96 | 4.15% |  |  |  |  |  |
| Zich et al., in prep | 0.11 | -0.30 | 0.48 | 15.82% |  |  |  |  |  |
| **Functional localizer** | **0.07** | **-0.14** | **0.27** | **46.54%** | **12.77** | **0.31** | **0.14** | **0.01** | **0.12** |
| Megumi et al., 2015 | 0.77 | 0.28 | 0.94 | 10.10% |  |  |  |  |  |
| Kim et al., 2015 | -0.50 | -0.94 | 0.59 | 6.56% |  |  |  |  |  |
| Kim et al., 2015 | -0.14 | -0.88 | 0.79 | 6.56% |  |  |  |  |  |
| Kirschner et al., 2018 | 0.65 | 0.31 | 0.84 | 13.39% |  |  |  |  |  |
| Kirschner et al., 2018 | 0.08 | -0.32 | 0.46 | 13.81% |  |  |  |  |  |
| Kirschner et al., in prep | 0.02 | -0.56 | 0.59 | 10.95% |  |  |  |  |  |
| Koush et al., 2013 | 0.79 | -0.16 | 0.98 | 6.56% |  |  |  |  |  |
| Koush_et al., 2015 | 0.32 | -0.55 | 0.85 | 8.31% |  |  |  |  |  |
| MacInnes et al., 2016 | -0.27 | -0.67 | 0.24 | 12.44% |  |  |  |  |  |
| Pamplona et al., in prep | 0.05 | -0.51 | 0.59 | 11.31% |  |  |  |  |  |
| **No-feedback run** | **0.24** | **-0.13** | **0.55** | **25.80%** | **23.05** | **0.01** | **0.61** | **0.15** | **0.38** |
| Marins et al., 2015 | -0.48 | -0.83 | 0.13 | 17.15% |  |  |  |  |  |
| McDonald et al., 2017 | -0.14 | -0.62 | 0.43 | 19.11% |  |  |  |  |  |
| McDonald et al., 2017 | 0.05 | -0.40 | 0.49 | 23.84% |  |  |  |  |  |
| Papoutsi et al., 2020 | -0.29 | -0.87 | 0.64 | 9.52% |  |  |  |  |  |
| Papoutsi et al., 2020 | -0.71 | -0.96 | 0.16 | 9.52% |  |  |  |  |  |
| Young et al., 2017 | 0.23 | -0.30 | 0.65 | 20.86% |  |  |  |  |  |
| **ROI-engaging run** | **-0.16** | **-0.51** | **0.23** | **27.66%** | **7.47** | **0.19** | **0.33** | **0.05** | **0.21** |

Table S6: **Averaged weighted Spearman correlations between pre-training activity levels and neurofeedback learning success during the first neurofeedback run, clustered by the paradigm of the pre-training run**
